# Chronic activation of corticospinal tract neurons after pyramidotomy injury enhances neither behavioral recovery nor axonal sprouting

**DOI:** 10.1101/2024.10.25.620314

**Authors:** Zimei Wang, Matthew Brannigan, Logan Friedrich, Murray G. Blackmore

## Abstract

Modulation of neural activity is a promising strategy to influence the growth of axons and improve behavioral recovery after damage to the central nervous system. The benefits of neuromodulation likely depend on optimization across multiple input parameters. Here we used a chemogenetic approach to achieve continuous, long-term elevation of neural activity in murine corticospinal tract (CST) neurons. To specifically target CST neurons, AAV2-retro-DIO-hM3Dq-mCherry or matched mCherry control was injected to the cervical spinal cord of adult Emx1-Cre transgenic mice. Pilot studies verified efficient transgene expression in CST neurons and effective elevation of neural activity as assessed by cFos immunohistochemistry. In subsequent experiments mice were administered either DIO-hM3Dq-mCherry or control DIO-mCherry, were pre-trained on a pellet retrieval task, and then received unilateral pyramidotomy injury to selectively ablate the right CST. Mice then received continual clozapine via drinking water and weekly testing on the pellet retrieval task, followed by cortical injection of a viral tracer to assess cross-midline sprouting by the spared CST. After sacrifice at eight weeks post-injury immunohistochemistry for cFos verified elevated CST activity in hM3Dq-treated animals and immunohistochemistry for PKC-gamma verified unilateral ablation of the CST in all animals. Despite the chronic elevation of CST activity, however, both groups showed similar levels of cross-midline CST sprouting and similar success in the pellet retrieval task. These data indicate that continuous, long-term elevation of activity that is targeted specifically to CST neurons does not affect compensatory sprouting or directed forelimb movements.

## INTRODUCTION

Damage to the central nervous system (CNS) leads to disruption of neural connectivity and loss of function. In principle, efforts to restore connectivity and function can focus either on the regeneration of axons that were directly damaged by injury or, alternatively, can aim to increase the number and/or efficacy of connections made by residual axons that survived the injury. In the case of spinal cord injury, the task of evoking regeneration from injured axons, especially long-distance growth, has proven challenging and slow to advance (Fawcett, 2020; Zheng and Tuszynski, 2023). At the same time there is growing awareness that even injuries classified as clinically complete often leave at least some spared axons within residual tissue that spans the injury (Chen et al., 2016; Friedli et al., 2015; Sherwood et al., 1992). These residual axons carry signals that are functionally sub-threshold, but which can be amplified for therapeutic benefit by providing additional stimulation to target fields (Angeli et al., 2014; Darrow et al., 2019; Harkema et al., 2011; Moritz et al., 2024; Rowald et al., 2022). Thus, although long-distance growth from injured axons will likely be needed for full recovery, approaches that aim to enhance collateralization and connectivity from spared axons are of high interest for their potential to offer tangible, albeit partial, improvements in function in the near term.

One of the most widely used pre-clinical models to assess new collateral growth from spared axons is unilateral pyramidotomy. This model involves a one-sided transection at the level of medullary pyramid that specifically targets the corticospinal tract (CST), an important mediator of sensory modulation and fine motor control (Fink and Cafferty, 2016; Kathe et al., 2014; Lemon, 2008; Starkey et al., 2005). Pyramidotomy deprives one side of the spinal cord of input from the CST, a predominantly crossed tract, and thus serves as a platform to test the ability of candidate treatments to enhance compensatory cross-midline sprouting and functional recovery from the intact CST. In addition to numerous compound and gene-based interventions, a promising means to enhanced compensatory CST sprouting is the use of neural activity as a pro-growth stimulus(Jara et al., 2020; Martin, 2016). Multiple studies employing electrical or transcranial magnetic stimulation of CST cell bodies in sensorimotor cortex have shown enhanced growth and collateralization from their spinal axons (Boato et al., 2023; Brus-Ramer et al., 2007; Carmel et al., 2013, 2010; Jack et al., 2018). Important questions remain, however, regarding the optimal use of neural stimulation. Because electrical and transcranial magnetic techniques broadly affect neurons and axons of passage in the stimulation field, they cannot determine the extent to which growth is stimulated through direct effects on CST neurons. Thus, it is not clear whether more targeted stimulation might be more effective than whole-cortex approaches. In addition, practical considerations typically limit the stimulation that each subject can receive, raising the question of whether stimulation that is more continual across the day or extended into more chronic time-points after injury might also be more effective.

The development of chemogenetic approaches, in which neurons are rendered sensitive to otherwise inert ligands by forced expression of exogenous, activity-modulating receptors, offers new opportunities to address both questions. Elevation of neural activity can be restricted to only a genetically defined subset of cells that express the exogenous receptor, while the administration of activating ligands via animal injection or drinking water offers a practical means to achieve long term activation across cohorts of animals. Here we used a chemogenetic approach to test the effects of continual, chronic elevation of CST activity in a pyramidotomy model of unilateral axon damage. Activating Gq-DREADD constructs were targeted broadly and specifically to the CST population using an intersectional transgenic/viral approach, and long-term elevation of activity was maintained by ligands supplied in drinking water. Immunohistochemistry for cFos confirmed long-term elevation of activity in CST neurons after pyramidotomy. Despite this increased activity, however, animals showed no evidence for either enhanced or suppressed sprouting into denervated spinal tissue, nor improvements in forelimb function as assessed by pellet retrieval. These results establish experimental parameters for specific, long-term stimulation of CST neurons and provide an important point of reference for the larger effort to optimize neural stimulation therapies.

## MATERIALS AND METHODS

### Animals

Experiments used male and female mice aged 7-8 weeks at the start of experiments, obtained by homozygous breeding from Emx1-Cre founders (C57/Bl6 background) obtained from Jackson Labs (#005628, RRID:IMSR_JAX:005628). Seventeen animals (eleven females, six males) were used for validation experiments to establish CST targeting and activation. Twenty-five animals (12 male, 13 female) were used for subsequent behavioral and histological experiments; all animals were randomly assigned to groups prior to the initiation of experiments. Mice were group housed under a 12/12 light/dark cycle (6am on time) with experiments carried out during the light phase. Animals had ad libitum access to food except during periods of behavioral testing (see below). All experiments were approved by the Institutional Animal Care and Use Safety Committee at Marquette University and in accordance with the National Institutes of Health Guidelines for the Care and Use of Laboratory Animals.

### Viral delivery

Viral delivery to the spinal cord followed the procedures of (Beine et al., 2022; Wang et al., 2022). Briefly, mice were anesthetized by a mixture of ketamine and xylazine (100 mg/kg and 10 mg/kg, respectively), an incision was made to expose the spinal cord between C2 and T2, and mice were mounted on a custom spine stabilization device. Viral particles were delivered to C5 spinal cord using a Hamilton syringe fitted with a pulled glass pipette at a rate of 0.05 μl/min to coordinates 0.3mm to the left of the midline and 0.8mm depth. After 0.5µl of viral particles were dispensed at this location the needle was raised 0.2mm and an additional 0.5µl were dispensed. Injected particles were either AAV2-Retro-hSyn-DIO-hM3D(Gq)-mCherry (Addgene #44361, 7.0 × 10^12^ GC/mL) or AAV2-Retro-hSyn-DIO-mCherry (Addgene #50459, 7.0 × 10^12^ GC/mL). After two additional minutes to minimize backflow the needle was retracted.

Viral delivery to the cortex followed procedures of (Kramer et al., 2021; Z. Wang et al., 2015). After anesthesia with a ketamine/xylazine cocktail (100mg/kg, 10mg/kg) animals received injections of AAV9-luc-EGFP into motor cortex through a pulled glass pipette fitted onto a 10μl Hamilton syringe at two locations: (relative to Bregma) anterior 0.0mm, left 1.3mm, depth 0.5mm and anterior 0.5mm, left 1.3mm, depth 0.5mm. At each site 0.5μl was delivered at a rate of .05μl/min using a Stoelting QSI pump (#53311). Following each injection the pipette was left in place for two minutes to minimize back flow. Viruses were designed in house and generated from the University of North Carolina Vector Core as previously described (Wang et al., 2018).

### Pyramidotomy

Following the procedures of (Zimei Wang et al., 2015) mice were anesthetized with a mixture of ketamine and xylazine (100 mg/kg and 10 mg/kg, respectively). Using a ventral approach the medullary pyramids were exposed by removal of the ventrocaudal region of the occipital bone with delicate rongeurs. The dura was punctured, and the left pyramid was cut completely using a micro feather scalpel. Mice were given a subcutaneous injection of Meloxicam (5 mg/kg) after the procedure. The mice’s general health and mobility were monitored daily throughout the post-injury survival period.

### CNO and Clozapine administration

Clozapine N-oxide (Abcam, ab141704) was dissolved in saline to a concentration of 1 mg/ml and injected intraperitoneally to a final dose of 5 mg/kg. The mice were collected 6 hours after injection (Wang et al., 2018). Clozapine (HB6129; HelloBio) was dissolved in water to a concentration of 1mg/ml (Hilton et al., 2022) and provided in the home cage as the sole source of drinking water. Clozapine water was refreshed every two days for the duration of its administration.

### Pellet retrieval task

Mice were motivated for the pellet retrieval task by food deprivation. This entailed complete food removal until 10% of body weight was lost and then daily access to food for a two-hour period with daily monitoring to ensure maintenance within 10% to 15% reduction from pre-restriction weight. For the pellet retrieval task animals were placed within a plexiglass enclosure with a 1cm slit in the front right corner, which allowed reaches only by the right forelimb. Millet food pellets were presented on a 3D-printed tray in which pellets rested on a pillar with 3mm diameter and 10mm height, located 1cm from the enclosure’s opening. In this configuration any attempt to drag the pellet into the enclosure resulted in the pellet falling out of reach; pellets could be retrieved only by grasping and lifting over the intervening space. Each pellet was presented for 30 seconds, with 80 total pellets offered in initial training sessions and 40 pellets presented in subsequent testing sessions. Task outcomes were scored by imaging and analyzing the presentation tray after each run. Pellets that remained on the pillars were scored as “missed”, pellets still visible on the tray but off the pillars were “displaced”, and pellets that were missing from the tray were “retrieved.” Pellet retrieval was expressed as a percent of total pellets presented and contacted pellets was calculated as pellets retrieved plus pellets displaced as a percent of total pellets presented. Animals were trained five days a week for two weeks. Tests of CNO or clozapine effects were performed for two days in a row with administered ligand, followed by a day of drug washout and no testing, and then two days of testing in the absence of ligand. Post-injury testing was conducted each week on two adjacent days and the reported scores reflecting the average of the two days.

### Tissue Processing and Immunohistochemistry

Animals were euthanized with CO_2_ followed by transcardiac perfusion with 4% paraformaldehyde (15710-Electron Microscopy Sciences) in Phosphate Buffered Saline (PBS, Sigma P4474). The brain and spinal column were postfixed overnight in 4% PFA, followed by fine dissection of brain, medulla and cervical spinal cord (a 6mm segment spanning C1-C6), which were stored in PBS. Cortex, spinal cord, and medullas were embedded in 12% gelatin (VWR, 71003-404) in PBS and cut via Vibratome (Leica VT1200) to yield 100μm sections. For cFos immunohistochemistry cortical sections were blocked in a 10% Normal Goal Serum (NGS, Gibco, 16210064), 3% Triton-X-100 (Sigma 9002-93-1) and then incubated overnight with anti-cFos antibody (1:400, Cell Signaling Technology, 2250, RRID:AB_2247211) in 3% NGS and 0.4% Triton-X. For PKCγ immunohistochemistry transverse sections of C3, C4, and C5 spinal cord were similarly blocked and incubated with anti-PKCγ antibody (1:500, Abcam, ab71558, RRID:AB_1281066). Sections were then incubated in goat anti-Rabbit IgG (H+L) Cross-Adsorbed Secondary Antibody, Alexa Fluor 647 (1: 500) (Invitrogen, #A-21244) for 2 h at room temperature in 3% NGS and 0.4% Triton-X for two hours, rinsed, and mounted on slides.

### Imaging and Data Analysis

For quantification of cFos signal, fluorescent images in two channels corresponding to mCherry and cFos signal were acquired at 20X using a Nikon A1R+ laser scanning confocal microscopy system on a Nikon Ti2-E inverted microscope. Using NIS Elements software individual CST cell bodies were manually outlined using signal in the mCherry channel to define a region of interest. Mean pixel intensity within this areas of interest was then extracted for the cFos channel and background-corrected by subtracting the mean pixel intensity in an adjacent, mCherry-negative region. Values were collected for at least 60 individual cells in each of two replicate tissue sections from all animals. Thresholds for assigning a cell as positive for cFos were calculated as the average background values plus three times the standard deviation of all background regions in that section. For quantification of PKCγ signal the right and left dorsal columns were manually outlined in 20X fluorescent images and the average pixel intensity determined for each. The mean pixel intensity in the right (injured) column was expressed as a percent of the value in the left (uninjured) column.

### Quantification of axon sprouting

Using a Nikon A1R+ laser scanning confocal microscopy system on a Nikon Ti2-E inverted microscope 20x images were acquired of transverse sections of cervical spinal cord at levels C3, C4, and C5. Virtual lines were drawn at 400μm to the left of the midline (uninjured side) and at 200 μm, 400 μm, and 600 μm to the right of midline (denervated side), with the width of the line set to 10 μm. The total number of contacts between EGFP+ CST axons and each line was manually counted by two independent, blinded observers and then averaged. The total number of EGFP+ axons in the medulla was determined by imaging the medullary region at 60X. The number of axonal profiles within three sampling regions (squares, 100 μm on each side) was averaged to calculate axon density and then multiplied by the total cross-sectional area of the medulla to calculate the total number of axons. Axon counts from the spinal cord were divided by the total number of axons detected in the medulla of the same animal to yield axon index values.

### Statistical analyses

All surgeries and quantification were performed by blinded personnel. Behavioral outcomes were assessed for significance using two-way ANOVA with Sidak’s post-tests to compare groups in the presence and absence of DREADD-activating ligands and by repeated measure two-way ANOVA with Tukey’s multiple comparisons to determine group differences at multiple time points post-injury. Axon growth was compared by two-way ANOVA with Tukey’s multiple comparisons. Differences in mean cFos intensity were tested for significance using ANOVA with post-hoc Dunnett’s. All statistical tests were carried out using Graphpad 10.3.1.

## RESULTS

### Establishing experimental parameters from chronic DREADD-mediated activation of CST neurons

To express activating DREADD receptors (Alexander et al., 2009) selectively in CST neurons we delivered AAV2-Retro-hSyn-DIO-hM3D(Gq)-mCherry (hereafter Gq-DREADD) to the cervical spinal cord of adult Emx1-Cre mice, with AAV2-Retro-hSyn-DIO-mCherry as control (**Fig. 1A**). Animals were perfused three weeks later, with half of the animals receiving activating CNO ligand by intraperitoneal (IP) injection six hours prior to sacrifice. Cortices were then examined for evidence of retrograde transduction (mCherry expression) and neural activity (cFos immunohistochemistry). As expected, mCherry signal was readily detectable in layer V cortex in all groups, indicating expression of mCherry control or mCherry-hM3D(Gq) protein in CST neurons (**Fig. 1B-D**). cFos signal was detected in fewer than five percent of CST neurons in mCherry control animals, regardless of CNO injection, and was similarly low in animals treated with Gq-DREADD but which received no activating CNO (**Fig. 1B, C, G**). In contrast, cFos was detected in more than 70% of mCherry+ CST cell bodies in Gq-DREADD-transduced animals stimulated with CNO (**Fig. 1D**). Quantification confirmed a large and significant increase in average pixel intensity of cFos signal within mCherry+ CST bodies in Gq-DREADD animals compared to both mCherry and Gq-DREADD/no CNO controls (p<.001, ANOVA with post-hoc Dunnett’s). These data confirm the ability of an AAV2-Retro/transgenic intersectional strategy to selectively increase neural activity in CST neurons.

**Figure 1.**
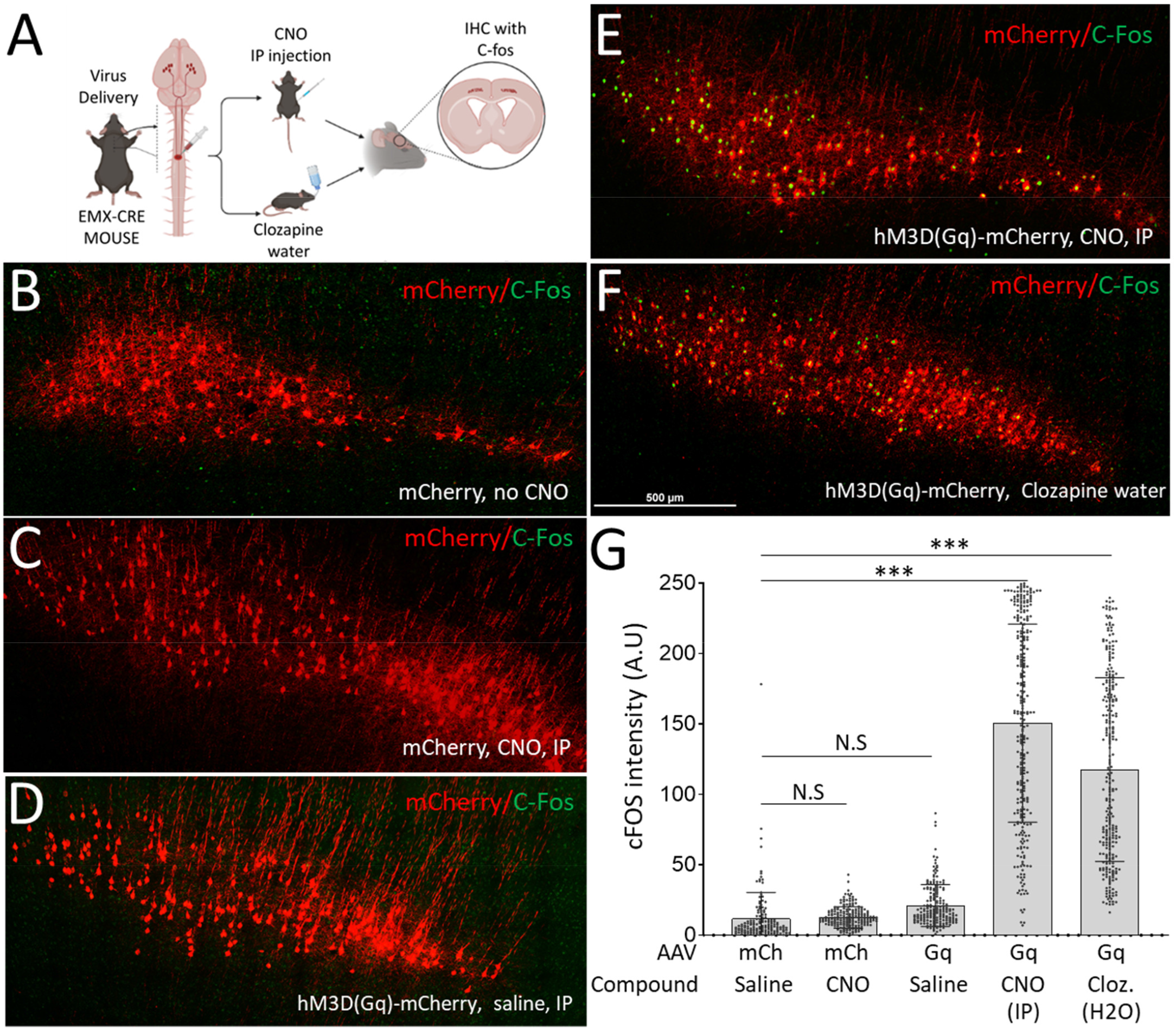
DREADD-mediated activation of CST neurons. (A) An experimental overview illustrates the retrograde delivery of cre-dependent Gq-DREADD-mCherry or mCherry control followed by administration of CNO by intraperitoneal injection (IP) or clozapine by drinking water. (B-F) show CST neurons identified by mCherry reporter (red) and cFOS immunohistochemistry in green. Control animal that received only mCherry AAV (B), mCherry AAV and CNO (C), or Gq-DREADD and no CNO (D) show minimal cFOS expression. In contrast, animals that received Gq-DREADD AAV and intraperitoneal CNO (E) or clozapine in drinking water (F) show extensive cFOS expression. (G) Quantification of pixel intensity of cFOS signal within mCherry-expressing CST neurons confirms significant upregulation in AAV-Gq-DREADD animals that received stimulating ligand either by IP injection or through drinking water. Error bar shows SEM. N > 100 individual cells three 3 separate animals per treatment, *** p<.001, one way ANOVA with post-hoc Dunnett’s.

IP delivery of CNO yields neuronal activation that likely lasts only several hours (Wang et al., 2018). We therefore next explored delivery of a DREADD activator in drinking water as a more practical means to achieve long-term CST activation. As previously, adult Emx1-Cre mice received AAV2-Retro injection of Cre-dependent Gq-DREADD-mCherry or mCherry control to cervical spinal cord. Three weeks later mice were supplied for three days with drinking water that contained clozapine, the active metabolite of CNO, and then perfused (Gomez et al., 2017). As in the prior experiment, cFos was detected in just a few percent of CST neurons in mCherry control animals but was visible in more than 60% of CST neurons in Gq-DREADD animals (**Fig. 1E, F**). Quantification again confirmed a large and significant increase in cFos signal in CST cell bodies in Gq-DREADD animals compared to mCherry control (**Fig. 1G**). Thus, retrograde transduction of CST neurons with DREADD receptors, in conjunction with continual supply of clozapine ligand in drinking water, stably elevates CST activity.

### Effect of chronic activation of CST neurons on forelimb recovery after pyramidotomy injury

We next asked how elevation of CST neural activity affected forelimb function. As previously, groups of adult Emx1-Cre transgenic mice received cervical injection of AAV2-Retro-hSyn-DIO-hM3D(Gq)-mCherry or AAV2-Retro-hSyn-DIO-mCherry control. After a week to recover from surgery, mice were pre-trained on a pellet retrieval task in which animals reached through a narrow slit with the right forelimb to attempt retrieval of a food reward (**Fig. 2A**). Food pellets were placed atop 3D-printed pillars located 1cm from the slit. Thus, animals were obligated to lift the pellet over the intervening space, favoring grasp of the pellet and preventing scooping or dragging solutions. During the initial training period animals were exposed to eighty pellets per session and were subsequently tested with forty pellets. The number of pellets successfully moved into the retrieval cage and the number of pellets that were contacted (pellets eaten plus pellets knocked from the pillar were determined for each animal and expressed as a percent of total pellets presented.

**Figure 2.**
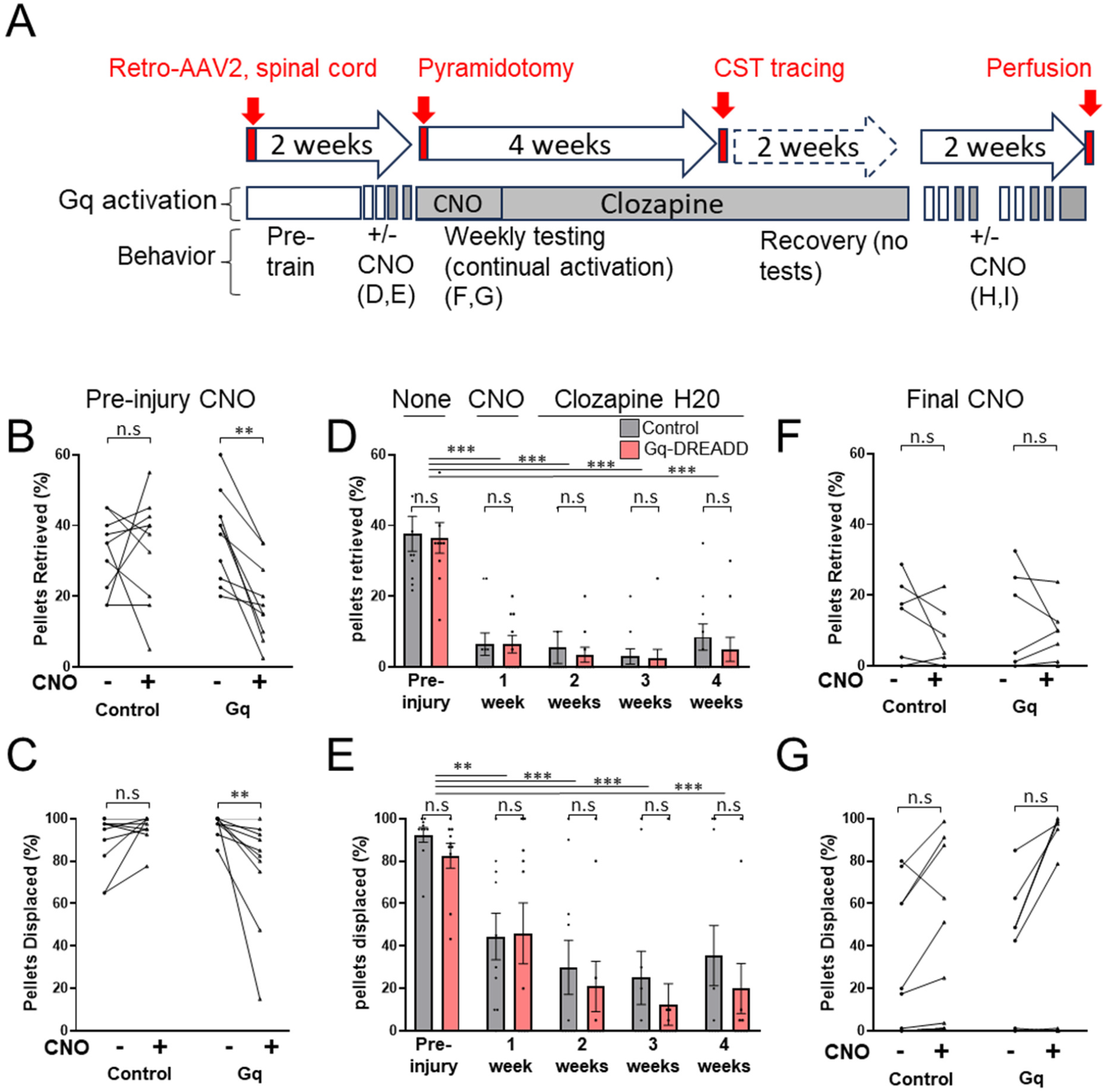
Chronic elevation of CST activity does not improve forelimb function after pyramidotomy injury. (A) Experimental sequence of delivery of Gq-DREADD or control mCherry followed by training, injury, and chronic activation. (B, C) Prior to injury, acute CNO delivery reduces the success of pellet retrieval (B) and pellet contact (C) in Gq-treated but not control animals. (D,E) Pyramidotomy injury significantly impairs pellet retrieval (D) and contact (E), but chronic DREADD activation has no effect on pellet retrieval or displacement compared to control-treated animals. (F,G) Acute DREADD activation eight weeks after injury produces no significant change in pellet retrieval performance. N=10 animals per group. Error bars represent SEM. B,C,F,G : **p < 0.01, two-way ANOVA with Sidak’s post-tests. D,E: ***p < 0.01, two-way RM ANOVA with Tukey’s multiple comparisons.

After two weeks of training, both control and Gq-DREADD animals contacted an average of more than 90% of pellets and reached stable retrieval rates that averaged 30.3% (±3.8% SEM) and 37.3% (±3.3% SEM), respectively. To determine how DREADD-mediated activation affected task performance prior to injury, all animals received injections of CNO fifteen minutes prior to testing on the final two days. Average performance across CNO-treated days was then compared to the pre-CNO baseline. In control animals CNO injection produced no significant effect on retrieval success (30.3% ±3.8% SEM versus 29.3% ±5.7% SEM, p=0.87, paired t-test) or pellet contact (92.0% ±2.7% SEM versus 86.3% ±9.3% SEM, p=0.45, paired t-test) (**Fig. 2D, E**). Thus, neither CNO nor the stress of IP injections affected task performance in control animals. In contrast, injection of CNO to Gq-DREADD animals resulted in significant reduction in both average retrieval (18.5% ±3.5% SEM vs. 37.3% ±3.3% SEM baseline, p=.001, paired t-test) and contact (76.3% ±8.2% SEM vs. 97.0±1.5% SEM baseline, p=.001, paired t-test) (**Fig. 2D, E**). These data show that DREADD activation of CST neurons just prior to task performance may interfere with fine forelimb function in animals that learned the pellet retrieval task in the absence of exogenous CST activation.

Animals then received unilateral pyramidotomy injury to sever CST axons that normally innervate the right spinal cord. In the first week after injury, when animal’s drinking may have been disrupted, we delivered CNO twice daily by IP injection (see **Fig. 2A** for experimental timeline). Starting one week after injury and continuing until six weeks post-injury animals received clozapine via drinking water. Animals were tested weekly on the pellet retrieval task to monitor the effects of injury and CST activation on forelimb function. Pyramidotomy injury strongly reduced task performance in both control and Gq-DREADD animals, with only 7.3%±4.2% SEM and 8.25%±3.4% SEM retrieval rates, respectively, in the week following injury (**Fig. 2F**). The rate of pellet contact also declined after injury to an average of 38.5%±10.9% SEM and 44%±13.7% SEM in control and Gq-DREADD groups, respectively. Thus, animals mostly lost the ability to grasp and retrieve pellets but retained some ability to contact and displace pellets from stands. In the four weeks after injury animals displayed no spontaneous recovery of task performance, with both groups averaging fewer than 10% of pellets retrieved in all testing periods (p>0.05 versus the first week post-injury, repeated measures 2-way ANOVA with Tukey’s multiple comparison) (**Fig. 2F, G**). Importantly, the Gq-DREADD group performed similarly to controls in both pellet retrieval and displacement at all timepoints, indicating that chronic CST activation had neither positive nor negative effects on forelimb function (p>.05 Gq-DREADD vs. Control at all timepoints, repeated measures 2-way ANOVA with Tukey’s multiple comparisons) (**Fig. 2F, G**). Overall, these data indicate that unilateral pyramidotomy injury produced a strong and lasting reduction in the execution of a grasp-dependent retrieval task, but that chronic chemogenetic activation of CST neurons had no effect on functional recovery.

Six weeks after injury, animals received cortical injections to enable tracing of CST axons (see below) and were not tested during a two-week recovery period. DREADD activation was continual through these two weeks, first via CNO injections and then with clozapine in drinking water. In weeks seven and eight after injury animals were tested on the pellet retrieval task four days per week, with the first two tests each week performed in conditions of maintained DREADD activation and the next two tests performed after at least 48 hours of drug washout (no injections, no clozapine in drinking water). As in the initial four weeks of testing, Gq-DREADD and control groups performed similarly during all sessions, indicating that chronic CST activation did not impact task performance. Interestingly, unlike the pre-injury results, Gq-DREADD animals performed identically in the presence or absence of activating ligand. Overall, chemogenetic activation of CST neurons had no effect on forelimb function in the pellet retrieval task after pyramidotomy injury.

Finally, to confirm the efficacy of chemogenetic activation in this cohort, prior to perfusion all animals were administered clozapine via drinking water for two full days. As previously, cortical sections were prepared and subjected to cFos immunohistochemistry. As expected, mCherry signal was abundant in layer V cortex of both control and Gq-DREADD treated animals, verifying successful retrograde transduction in all animals (**Fig. 3A, B**). In addition, Gq-DREADD animals showed significant elevation of c-Fos signal compared to control when quantified either as the average cFos signal intensity in CST (mCherry+) neurons (**Fig. 3C**) or as the percent of CST neurons in each animal with detectable cFos signal (**Fig. 3D**). These data support the prior validation data and confirm within this animal cohort that the combined viral / clozapine strategy resulted in effective retrograde transduction and chemogenetic elevation of CST activity.

**Figure 3.**
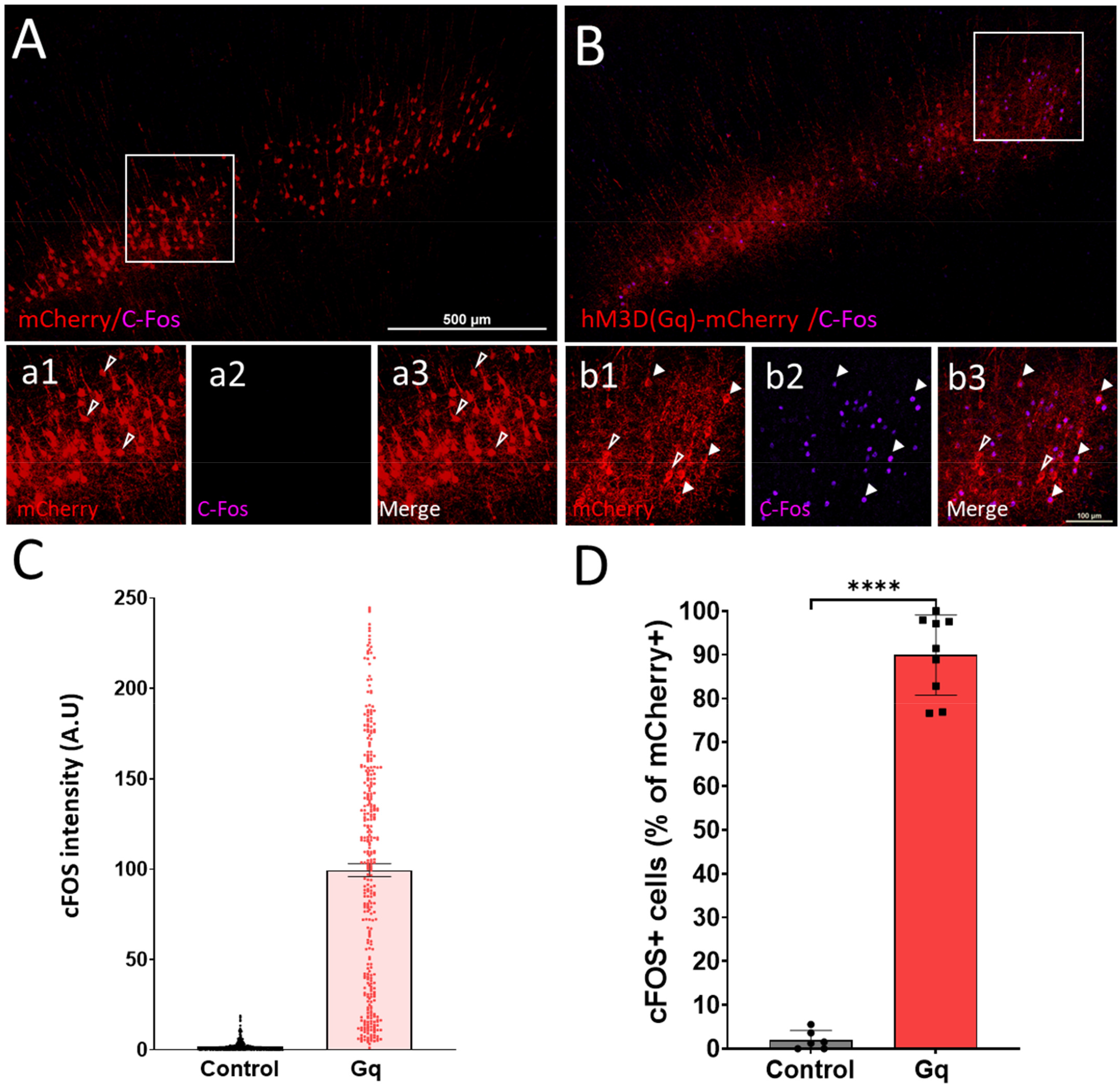
Confirmation of elevated neural activity in chronically stimulated CST neurons. Data were gathered from the same cohort of animals described in Figure 2 after being returned to clozapine delivered by drinking water for two full days. (A, B) Retrogradely transduced CST neurons (cherry+, red) show minimal cFOS expression in animals treated with DiO-mCherry control (A) but strong upregulation in animals treated with DiO-Gq-DREADD-mCherry (B). (C) Quantification of cFOS intensity shows an increase in signal in Gq-DREADD expressiong CST neurons compared to control; each dot is an individual cell. (D) An alternative quantification of the percent of CST neurons that express detectable cFOS also shows a significant increase in Gq-treated animal; each dot represents the percent of cFOS-expressing CST neurons in an individual animal. N=429 cells (control, C), 343 cells (Gq, C), 6 animals (control, D), and 9 animal (Gq, D). Error bars showSEM. ***p<.001, unpaired t-test.

### Cross-midline CST sprouting after unilateral pyramidotomy is unaffected by chronic activation

We examined spinal tissue from the set of behaviorally tested animals (above) to determine whether chemogenetic activation affected CST axon growth. We first verified completeness of the unilateral pyramidotomy injuries using transverse sections of cervical spinal cord and immunohistochemistry for PKCγ, a protein associated with CST axon tracts. In all animals from both control and Gq-DREADD groups PKCγ signal was prominent in the left dorsal column, the location of the intact CST, but abolished in the right dorsal column (**Fig. 4A, B**). Images of spinal cords from all animals are available in Supplemental Figure 1. Quantification of the relative right/left PKCγ intensity showed a similar loss signal in the right CST in both groups. These data confirm that both control and Gq-DREADD groups received effective pyramidotomy injuries.

**Figure 4.**
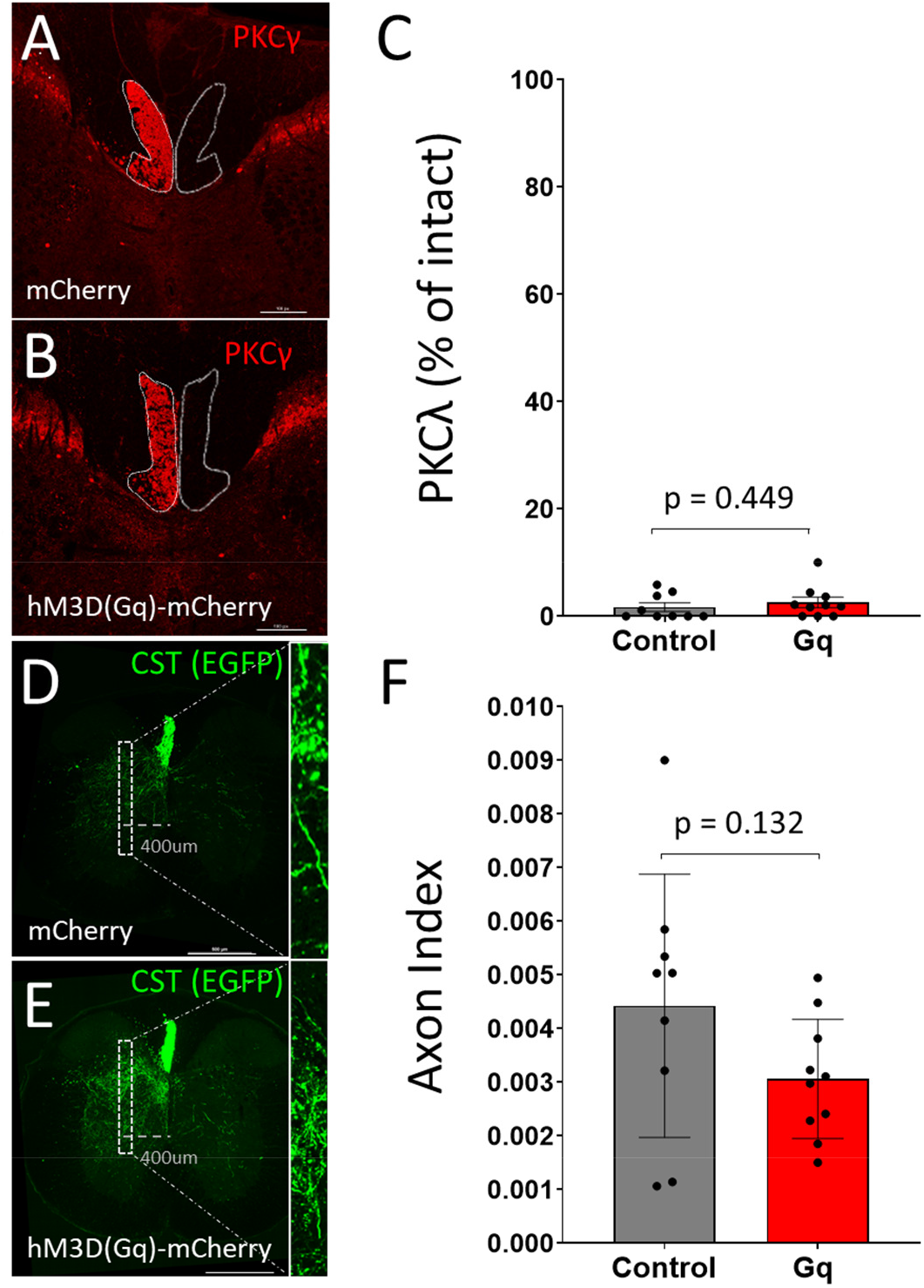
Control and Gq-DREADD treated animals were similarly injured and displayed similar CST collateralization in the intact side of the spinal cord. (A, B) show transverse sections of cervical spinal cord with bright PKC gamma immunoreactivity in the left dorsal CST (white outline) but no signal in the contralateral (injured) CST. (C) Quantification of total PKC gamma immunofluorescence in the right dorsal CST, expressed as a percent of the left, confirms strong reduction in both animal groups. (D, E) show transverse sections of cervical spinal cord with CST axons labeled by EGFP (green). White boxes indicate sampling regions to quantify the number of CST collaterals located 400µm to the left of the midline. (F) Quantification of CST collateralization into left (intact) spinal cord detects no significant difference between groups. Error bars show SEM. P>.05, unpaired t-test. Scale bar = 100µm (A-B), 500µm (C-D).

We next measured the ramification of CST collaterals in cervical spinal cord using AAV9-EGFP tracer that had been injected to the right cortex four weeks prior to sacrifice (see **Fig. 2C** for experimental timeline). As expected, in transverse sections of cervical spinal cord EGFP+ CST axons were visible as a tight bundle of axons in the left (intact) dorsal column and as tortuous collaterals extending mostly into the left grey matter (**Fig. 4D, E**). We first tested whether DREADD activation affected CST ramification in the left (intact) grey matter by quantifying intersections between EGFP+ profiles and a virtual line placed 400µm to the left of the midline (**Fig. 4D, E**). To account for potential variability in viral labeling these values were normalized to total EGFP+ profiles in the medullary pyramids (axon index). Control and Gq-DREADD animals showed similar axon indexes (**Fig. 4F;** p>.05, t-test), indicating that chronically elevated CST activity did not alter collateral density in intact spinal tissue.

We next assessed cross-midline sprouting of the left CST into right grey matter. Transverse sections of C3, C4, and C5 spinal cord were prepared and EGFP+ profiles were quantified at distances of 200µm, 400µm, and 600µm to the right of the midline (**Fig. 5 A-F**). As previously, axon counts were normalized to the number of EGFP+ profiles detected in the medullary pyramids (**Fig. 5G, H**). Quantification of total cross-midline sprouting across all cervical levels showed no significant difference between groups at any distance across the midline (**Fig. 5I)**. Similarly, comparisons made within each of the three cervical levels also revealed no significant differences **(Supplemental Figure 1**). Images of all spinal sections from all animals are available in **Supplemental Figure 1**. These data show that prolonged chemogenetic activation of CST neurons after unilateral pyramidotomy, the completeness of which was confirmed by terminal immunohistochemistry (see **Fig. 3**, above), did not affect cross-midline growth by CST neurons.

**Figure 5.**
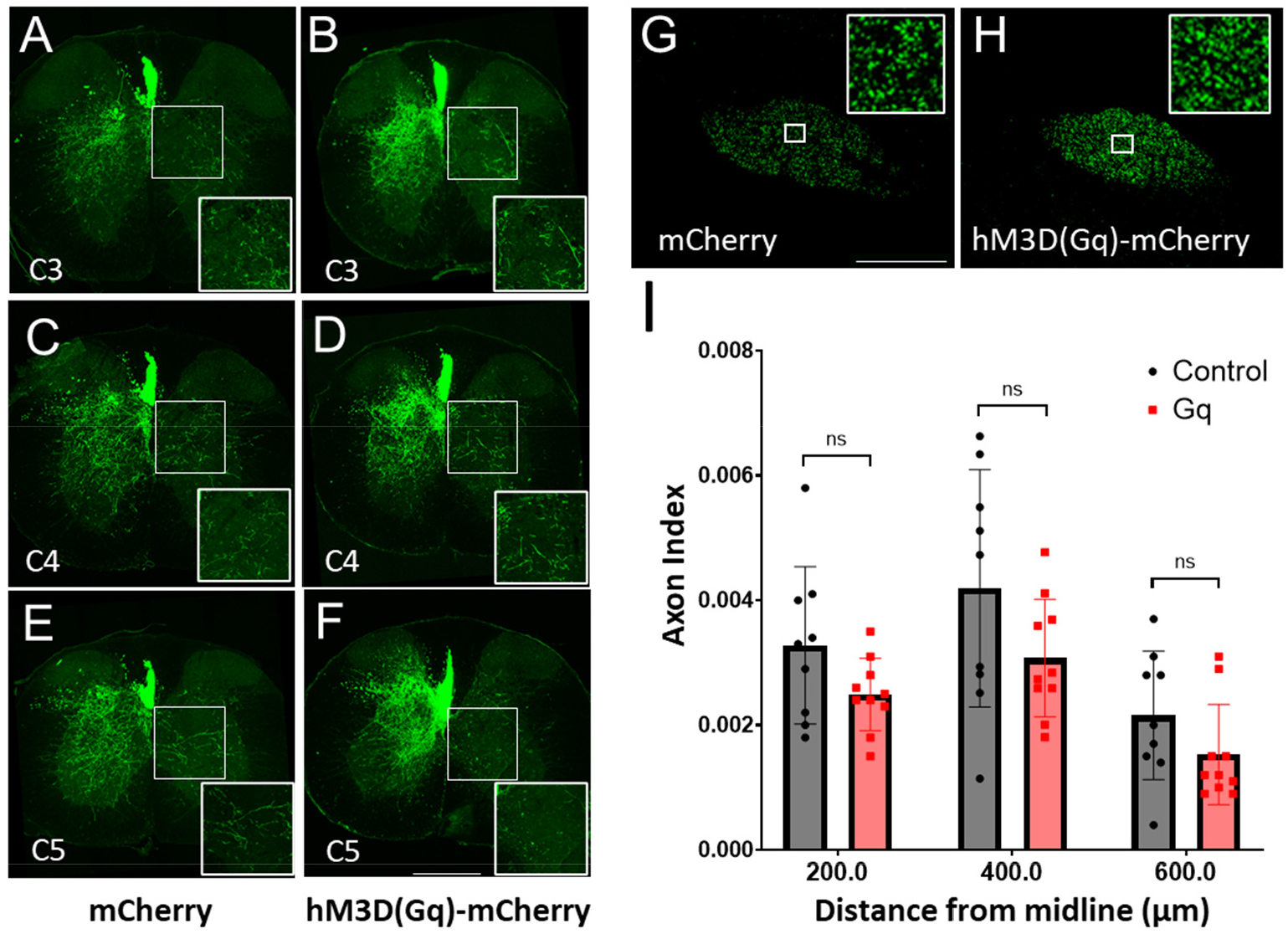
Elevation of CST activity has no effect on cross-midline sprouting after pyramidotomy injury. (A-F) show spinal cord sections at level C3 (A,B), C4 (C,D), or C5 (E,F), eight weeks after left pyramidotomy. CST axons (green) were labeled by injection of EGFP to the right cortex. Animals that received control AAV-mCherry (A, C, E) and AAV-Gq (B,D, F) show similar amounts of cross-midline CST axon growth. (G, H) show examples of coronal sections of the medulla with similar numbers of CST axons (green), illustrating comparable transduction. (I) Quantification of cross-midline CST axon growth, normalized to the number of virally transduced CST axons detected in the medulla, detects no significant difference between mCherry and Gq-treated animals. Scale bars are 500µm (A-F) and 200µm (G,H). ns indicates p>.05, 2-way ANOVA with Tukey’s post hoc test. N=9 (control), and N=10 (Gq). Error bars show ±SEM.

## DISCUSSION

This study used genetically targeted DREADD constructs to explore the effects of cell-specific, chronic elevation of neural activity on axon sprouting and behavioral outcomes after CNS damage. The approach succeeded in specific and widespread activation of CST neurons, as evidenced by elevated cFos expression and the observation that acute, first-time DREADD activation altered behavioral outcomes in a CST-depending forelimb task. Chronic elevation of CST activity did not, however, increase axon sprouting in the spinal cord or promote recovery of pellet retrieval. This outcome adds to a collection of studies that have employed various means to manipulate neural activity, and which have observed variable effects on axon growth and recovery. These divergent outcomes may provide an opportunity to guide optimization efforts by comparing input parameters and efficacy across studies. Below we provide a brief survey of prior reports that provide context for the current findings and then examine key treatment parameters that may help explain the differential success.

Numerous studies from multiple labs indicate that when stimulation is applied in a non-selective manner to whole cortical tissue it generally enhances both CST sprouting in the spinal cord and improves the execution of forelimb tasks. Specifically, broad cortical stimulation by electrical or transcranial magnetic approaches have shown benefits in various models of rodent injury, including pyramidotomy models like the one used here (Batty et al., 2020; Boato et al., 2023; Carmel et al., 2013, 2010; Jack et al., 2018; Yang et al., 2019). Interestingly, although effects are positive at the group level, considerable variability is observed between animals. For example, even within a single cohort performed by the same experimenters and with presumably consistent procedures, the effect of cortical stimulation on total cross-midline growth in pyramidotomy models can range from more than ten times the average of unstimulated controls in some animals to no increase in others (Carmel et al., 2013, 2010). In aggregate, results from electrical and transcranial magnetic studies strongly support the potential for positive growth and behavioral effects but also highlight the sensitivity of outcomes to input variables that may be difficult to standardize.

Results from optogenetic and chemogenetic approaches have also frequently been positive, although not universally so. For example, optogenetic stimulation of layer V cortical neurons using a Thy1-ChR2 transgenic line increased cross-midline sprouting of CST neurons more than two-fold after spinal hemisection (Wu et al., 2022), and optogenetic stimulation of virally transduced CST neurons similarly increased total axon length in the spinal cord of uninjured animals (Yang and Martin, 2023) and in animals after stroke (Wahl et al., 2017). Regarding chemogenetic approaches, prior work found that stimulation of CST neurons increased spinal axon growth in uninjured animals (Yang and Martin, 2023) and in animals after stroke (Yang et al., 2024), and elevated the number of Vlgut1 puncta, a surrogate for CST axon density, after spinal hemisection (Lin et al., 2023). On the other hand, other studies have reported small or no positive effects of opto- or chemogenetic stimulation of cortex. For example, in one study optogenetic stimulation of layer V neurons after spinal injury produced only a minor sprouting effect within several hundred microns of the injury site and no gains in forelimb function (Mesquida-Veny et al., 2022). Similarly, chemogenetic stimulation of CST neurons had no effect on axons density or behavioral recovery after dorsal spinal hemisection (Van Steenbergen et al., 2023), nor on pellet retrieval success (Lin et al., 2023). These results are in line with the current demonstration that CST activity was definitively elevated without producing any enhancement of axon growth or behavior. Thus, it appears that differences in approach with genetic or virally targeted stimulation strategies can lead to highly divergent outcomes, highlighting the need to identify and optimize the key input variables.

We have considered four non-exclusive explanations for differential success. The first is that cell-specific stimulation may be less effective than whole-tissue approaches. In the current study we used an intersectional viral approach to achieve activation that was highly selective to CST neurons, a key distinction from electrical stimulation paradigms and from non-selective cortical delivery of chemo- or optogenetic constructs. When delivered non-specifically, stimulation may spur axon growth in part through cell-extrinsic mechanisms (e.g. tissue-level rise in neurotrophins). If so, such benefits would be largely absent in conditions of selective activation like those used here. Consistent with this, it is notable that whereas non-selective stimulation techniques have yielded nearly universal efficacy across labs, stimulation targeted specifically to CST neurons by intersection viral means has yielded mixed reports of both positive (Wahl et al., 2017) and small or neutral effects on axon growth and behavior (Lin et al., 2023; Yang et al., 2024). Another illustrative example comes from comparing electrical stimulation that was targeted specifically to CST axons in the medullary pyramids versus broadly to cortical tissue. The more selective stimulation increased axon growth by approximately 25% (Brus-Ramer et al., 2007), whereas stimulation delivered broadly to the cortex increased growth more than four-fold (Carmel et al., 2010). Importantly, these data come from the same pyramidotomy injury paradigm and from the same lab group. This is not to suggest that cell-specific stimulation is necessarily ineffective, as clear examples exist to support cell-autonomous effects of stimulation (Wahl et al., 2017; Yang and Martin, 2023). Nevertheless, the lack of effect in our paradigm of CST-selective activation adds to an emerging pattern that the effects of cell-type specific activation may be generally smaller than whole-tissue stimulation.

A second consideration regards the timing of the onset of activation after injury. In one recent report, chemogenetic stimulation of CST neurons initiated one day after spinal injury, but not seven days post-injury, not only failed to enhance growth but caused a significant reduction compared to control (Van Steenbergen et al., 2023). In our experiments we also initiated stimulation within a day of injury, and it is interesting to note that CST axon counts in stimulated animals showed a trend toward reduction, albeit non-significant (Figure 5). The idea that elevated neural activity could have negative effects in the immediate aftermath of injury is reminiscent of reports that rehabilitative training can interfere with recovery when initiated too soon after injury (Torres-Espín et al., 2018). Thus, our findings are consistent with the notion that neural stimulation may be less effective when applied immediately after injury.

Third, the lack of effect in our approach may reflect the frequency and/or duration of stimulation. It is well established that the efficacy of neuromodulation varies according to the specifics of the stimulation parameters (Jara et al., 2020). For example, theta burst stimulation is favored for enhancement of axon growth and behavioral recovery (Yang and Martin, 2023). Interestingly, another study found that high-frequency stimulation after spinal injury enhanced axon growth and behavioral recovery but that lower frequency stimulation had no behavioral benefit and worsened axonal dieback (Jack et al., 2018). In the present study we verified that chemogenetic stimulation led to an increase in overall activity, as verified by cFos expression, but we neither controlled more monitored the firing rate. It is therefore possible that the resulting activity was sub-optimal in frequency or other firing parameters. Along these same lines, a recent study showed that optogenetic stimulation of the cortex using a controlled theta burst paradigm enhanced CST axon growth into the ventral spinal cord but that chemogenetic activation did not (Yang and Martin, 2023). Thus, despite the practical advantages of chemogenetics for long term elevation of activity, the inability to optimize the firing pattern may lessen the net efficacy of this strategy.

Finally, a fourth distinguishing feature of our approach is that it broadly affected CST neurons across both hemispheres. Previous quantification of Retro-AAV2 transduction efficiency in our hands has indicated transgene expression in more than 90% of CST neurons (Wang et al., 2022, 2018), and examination of mCherry expression (Figures 1 and 3) suggests a similarly high efficiency in the current experiments. In contrast, prior experiments that expressed DREADD constructs in CST neurons relied on focal injection to the cortex (Lin et al., 2023; Wahl et al., 2017; Yang et al., 2024). Thus, unlike the current approach that produced an increase in activity in most CST neurons, prior experiments selectively activated a localized subset of CST neurons. It is well established that differential neural activity can drive the competitive capture of synaptic territory by axon terminals (Friel et al., 2014; Friel and Martin, 2007); MARTIN et al., 2007). Thus, one speculative explanation for the lack of growth enhancement is that the present strategy created conditions in which most CST axons experienced a similar increase in activity, rather than creating an activity advantage in selected neurons.

To summarize, we conceptualize ongoing efforts to harness neural activity for therapeutic gain after CNS injury as an optimization exercise across multiple variables, which include stimulation modality, degree of cellular specificity, timing of onset, duration, episodic versus continual delivery, firing frequency, and more. We have explored a novel region of the available optimization space by establishing procedures for stimulation that is specific to CST neurons, acts broadly across the population, is initiated within a day of injury, and which allows long-term, stable elevation of activity. The approach succeeded in its goal of long-term elevation of CST activity, as verified by cFos activity, and in this sense provides a technical roadmap. The lack of growth or behavioral response with this combination of parameters provides important guidance, albeit negative, for continuing efforts to optimize the deployment of neural activity-based treatments for recovery from spinal injury.

## Acknowledgments

This work was supported by grants from NINDS (R01NS107807) and the Bryon Riesch Paralysis Foundation

## AUTHOR CONTRIBUTIONS

Conceptualization, M.B.; Methodology, M.B., Z.W, L.F.; Investigation, Z.W, M.Br., L.F.; Formal Analysis – M.B. Z.W.; Writing – Original Draft, M.B., Writing – Review & Editing, M.B., Z.W.; Funding Acquisition, M.B.; Visualization – M.B. and Z.W.; Supervision, M.B

## REFERENCES

Alexander, G.M., Rogan, S.C., Abbas, A.I., Armbruster, B.N., Pei, Y., Allen, J.A., Nonneman, R.J., Hartmann, J., Moy, S.S., Nicolelis, M.A., McNamara, J.O., Roth, B.L., 2009. Remote control of neuronal activity in transgenic mice expressing evolved G protein-coupled receptors. Neuron 63, 27–39. 10.1016/J.NEURON.2009.06.014

Angeli, C.A., Edgerton, V.R., Gerasimenko, Y.P., Harkema, S.J., 2014. Altering spinal cord excitability enables voluntary movements after chronic complete paralysis in humans. Brain 137, 1394–1409. 10.1093/BRAIN/AWU038

Batty, N.J., Torres-Espín, A., Vavrek, R., Raposo, P., Fouad, K., 2020. Single-session cortical electrical stimulation enhances the efficacy of rehabilitative motor training after spinal cord injury in rats. Exp Neurol 324. 10.1016/J.EXPNEUROL.2019.113136

Beine, Z., Wang, Z., Tsoulfas, P., Blackmore, M.G., 2022. Single nuclei analyses reveal transcriptional profiles and marker genes for diverse supraspinal populations. J Neurosci JN-RM-1197-22. 10.1523/JNEUROSCI.1197-22.2022

Boato, F., Guan, X., Zhu, Y., Ryu, Y., Voutounou, M., Rynne, C., Freschlin, C.R., Zumbo, P., Betel, D., Matho, K., Makarov, S.N., Wu, Z., Son, Y.J., Nummenmaa, A., Huang, J.Z., Edwards, D.J., Zhong, J., 2023. Activation of MAP2K signaling by genetic engineering or HF-rTMS promotes corticospinal axon sprouting and functional regeneration. Sci Transl Med 15. 10.1126/SCITRANSLMED.ABQ6885

Brus-Ramer, M., Carmel, J.B., Chakrabarty, S., Martin, J.H., 2007. Electrical Stimulation of Spared Corticospinal Axons Augments Connections with Ipsilateral Spinal Motor Circuits after Injury. The Journal of Neuroscience 27, 13793. 10.1523/JNEUROSCI.3489-07.2007

Carmel, J.B., Berrol, L.J., Brus-Ramer, M., Martin, J.H., 2010. Chronic Electrical Stimulation of the Intact Corticospinal System after Unilateral Injury Restores Skilled Locomotor Control and Promotes Spinal Axon Outgrowth. Journal of Neuroscience 30, 10918–10926. 10.1523/JNEUROSCI.1435-10.2010

Carmel, J.B., Kimura, H., Berrol, L.J., Martin, J.H., 2013. Motor cortex electrical stimulation promotes axon outgrowth to brain stem and spinal targets that control the forelimb impaired by unilateral corticospinal injury. European Journal of Neuroscience 37, 1090–1102. 10.1111/EJN.12119

Chen, Y., He, Y., DeVivo, M.J., 2016. Changing Demographics and Injury Profile of New Traumatic Spinal Cord Injuries in the United States, 1972–2014. Arch Phys Med Rehabil 97, 1610–1619. 10.1016/J.APMR.2016.03.017

Darrow, D., Balser, D., Netoff, T.I., Krassioukov, A., Phillips, A., Parr, A., Samadani, U., 2019. Epidural Spinal Cord Stimulation Facilitates Immediate Restoration of Dormant Motor and Autonomic Supraspinal Pathways after Chronic Neurologically Complete Spinal Cord Injury. J Neurotrauma 36, 2325–2336. 10.1089/NEU.2018.6006

Fawcett, J.W., 2020. The Struggle to Make CNS Axons Regenerate: Why Has It Been so Difficult? Neurochem Res 45, 144. 10.1007/S11064-019-02844-Y

Fink, K.L., Cafferty, W.B.J., 2016. Reorganization of Intact Descending Motor Circuits to Replace Lost Connections After Injury. 10.1007/s13311-016-0422-x

Friedli, L., Rosenzweig, E.S., Barraud, Q., Schubert, M., Dominici, N., Awai, L., Nielson, J.L., Musienko, P., Nout-Lomas, Y., Zhong, H., Zdunowski, S., Roy, R.R., Strand, S.C., Brand, R. van den, Havton, L.A., Beattie, M.S., Bresnahan, J.C., Bézard, E., Bloch, J., Edgerton, V.R., Ferguson, A.R., Curt, A., Tuszynski, M.H., Courtine, G., 2015. Pronounced species divergence in corticospinal tract reorganization and functional recovery after lateralized spinal cord injury favors primates. Sci Transl Med 7, 302ra134. 10.1126/SCITRANSLMED.AAC5811

Friel, K.M., Martin, J.H., 2007. Bilateral Activity-Dependent Interactions in the Developing Corticospinal System. Journal of Neuroscience 27, 11083–11090. 10.1523/JNEUROSCI.2814-07.2007

Gomez, J.L., Bonaventura, J., Lesniak, W., Mathews, W.B., Sysa-Shah, P., Rodriguez, L.A., Ellis, R.J., Richie, C.T., Harvey, B.K., Dannals, R.F., Pomper, M.G., Bonci, A., Michaelides, M., 2017. Chemogenetics revealed: DREADD occupancy and activation via converted clozapine. Science (1979) 357, 503–507. 10.1126/science.aan2475

Harkema, S., Gerasimenko, Y., Hodes, J., Burdick, J., Angeli, C., Chen, Y., Ferreira, C., Willhite, A., Rejc, E., Grossman, R.G., Edgerton, V.R., 2011. Effect of epidural stimulation of the lumbosacral spinal cord on voluntary movement, standing, and assisted stepping after motor complete paraplegia: a case study. Lancet 377, 1938–1947. 10.1016/S0140-6736(11)60547-3

Hilton, B.J., Husch, A., Schaffran, B., Lin, T. chen, Burnside, E.R., Dupraz, S., Schelski, M., Kim, J., Müller, J.A., Schoch, S., Imig, C., Brose, N., Bradke, F., 2022. An active vesicle priming machinery suppresses axon regeneration upon adult CNS injury. Neuron 110, 51-69.e7. 10.1016/j.neuron.2021.10.007

Jack, A.S., Hurd, C., Forero, J., Nataraj, A., Fenrich, K., Blesch, A., Fouad, K., 2018. Cortical electrical stimulation in female rats with a cervical spinal cord injury to promote axonal outgrowth. J Neurosci Res 96, 852– 862. 10.1002/jnr.24209

Jara, J.S., Agger, S., Hollis, E.R., 2020. Functional Electrical Stimulation and the Modulation of the Axon Regeneration Program. Front Cell Dev Biol 8. 10.3389/FCELL.2020.00736

Kathe, C., Hutson, T.H., Chen, Q., Shine, H.D., McMahon, S.B., Moon, L.D.F., 2014. Unilateral pyramidotomy of the corticospinal tract in rats for assessment of neuroplasticity-inducing therapies. Journal of Visualized Experiments e51843. 10.3791/51843

Kramer, A.A., Olson, G.M., Chakraborty, A., Blackmore, M.G., 2021. Promotion of corticospinal tract growth by KLF6 requires an injury stimulus and occurs within four weeks of treatment. Exp Neurol 339. 10.1016/j.expneurol.2021.113644

Lemon, R.N., 2008. Descending Pathways in Motor Control. Annu Rev Neurosci 31, 195–218. 10.1146/annurev.neuro.31.060407.125547

Lin, X., Wang, X., Zhang, Y., Chu, G., Liang, J., Zhang, B., Lu, Y., Steward, O., Luo, J., 2023. Synergistic effect of chemogenetic activation of corticospinal motoneurons and physical exercise in promoting functional recovery after spinal cord injury. Exp Neurol 370, 114549. 10.1016/J.EXPNEUROL.2023.114549

Martin, J., 2016. Harnessing neural activity to promote repair of the damaged corticospinal system after spinal cord injury. Neural Regen Res 11, 0. 10.4103/1673-5374.191199

Martin, J., Friel, K., Salimi, I., Chakrabarty, S., 2007. Activity- and use-dependent plasticity of the developing corticospinal system☆. Neurosci Biobehav Rev 31, 1125–1135. 10.1016/j.neubiorev.2007.04.017

Mesquida-Veny, F., Martínez-Torres, S., Del Río, J.A., Hervera, A., 2022. Genetic control of neuronal activity enhances axonal growth only on permissive substrates. Molecular Medicine 28, 1–16. 10.1186/S10020-022-00524-2/FIGURES/6

Moritz, C., Field-Fote, E.C., Tefertiller, C., van Nes, I., Trumbower, R., Kalsi-Ryan, S., Purcell, M., Janssen, T.W.J., Krassioukov, A., Morse, L.R., Zhao, K.D., Guest, J., Marino, R.J., Murray, L.M., Wecht, J.M., Rieger, M., Pradarelli, J., Turner, A., D’Amico, J., Squair, J.W., Courtine, G., 2024. Non-invasive spinal cord electrical stimulation for arm and hand function in chronic tetraplegia: a safety and efficacy trial. Nature Medicine 2024 30:5 30, 1276–1283. 10.1038/s41591-024-02940-9

Rowald, A., Komi, S., Demesmaeker, R., Baaklini, E., Hernandez-Charpak, S.D., Paoles, E., Montanaro, H., Cassara, A., Becce, F., Lloyd, B., Newton, T., Ravier, J., Kinany, N., D’Ercole, M., Paley, A., Hankov, N., Varescon, C., McCracken, L., Vat, M., Caban, M., Watrin, A., Jacquet, C., Bole-Feysot, L., Harte, C., Lorach, H., Galvez, A., Tschopp, M., Herrmann, N., Wacker, M., Geernaert, L., Fodor, I., Radevich, V., Van Den Keybus, K., Eberle, G., Pralong, E., Roulet, M., Ledoux, J.B., Fornari, E., Mandija, S., Mattera, L., Martuzzi, R., Nazarian, B., Benkler, S., Callegari, S., Greiner, N., Fuhrer, B., Froeling, M., Buse, N., Denison, T., Buschman, R., Wende, C., Ganty, D., Bakker, J., Delattre, V., Lambert, H., Minassian, K., van den Berg, C.A.T., Kavounoudias, A., Micera, S., Van De Ville, D., Barraud, Q., Kurt, E., Kuster, N., Neufeld, E., Capogrosso, M., Asboth, L., Wagner, F.B., Bloch, J., Courtine, G., 2022. Activity-dependent spinal cord neuromodulation rapidly restores trunk and leg motor functions after complete paralysis. Nat Med 28, 260–271. 10.1038/s41591-021-01663-5

Sherwood, A.M., Dimitrijevic, M.R., Barry McKay, W., 1992. Evidence of subclinical brain influence in clinically complete spinal cord injury: discomplete SCI. J Neurol Sci 110, 90–98. 10.1016/0022-510X(92)90014-C

Starkey, M.L., Barritt, A.W., Yip, P.K., Davies, M., Hamers, F.P.T., McMahon, S.B., Bradbury, E.J., 2005. Assessing behavioural function following a pyramidotomy lesion of the corticospinal tract in adult mice. Exp Neurol 195, 524–539. 10.1016/j.expneurol.2005.06.017

Torres-Espín, A., Beaudry, E., Fenrich, K., Fouad, K., 2018. Rehabilitative Training in Animal Models of Spinal Cord Injury. J Neurotrauma 35, 1970–1985. 10.1089/NEU.2018.5906

Van Steenbergen, V., Burattini, L., Trumpp, M., Fourneau, J., Aljović, A., Chahin, M., Oh, H., D’ambra, M., Bareyre, F.M., 2023. Coordinated neurostimulation promotes circuit rewiring and unlocks recovery after spinal cord injury. J Exp Med 220. 10.1084/JEM.20220615

Wahl, A.S., Büchler, U., Brändli, A., Brattoli, B., Musall, S., Kasper, H., Ineichen, B. V., Helmchen, F., Ommer, B., Schwab, M.E., 2017. Optogenetically stimulating intact rat corticospinal tract post-stroke restores motor control through regionalized functional circuit formation. Nature Communications 2017 8:1 8, 1–16. 10.1038/s41467-017-01090-6

Wang, Z., Maunze, B., Wang, Y., Tsoulfas, P., Blackmore, M.G., 2018. Global connectivity and function of descending spinal input revealed by 3D microscopy and retrograde transduction. The Journal of Neuroscience 1196–18. 10.1523/JNEUROSCI.1196-18.2018

Wang, Z., Reynolds, A., Kirry, A., Nienhaus, C., Blackmore, M.G., 2015. Overexpression of Sox11 Promotes Corticospinal Tract Regeneration after Spinal Injury While Interfering with Functional Recovery. Journal of Neuroscience 35, 3139–3145. 10.1523/JNEUROSCI.2832-14.2015

Wang, Z., Romanski, A., Mehra, V., Wang, Y., Brannigan, M., Campbell, B.C., Petsko, G.A., Tsoulfas, P., Blackmore, M.G., 2022. Brain-wide analysis of the supraspinal connectome reveals anatomical correlates to functional recovery after spinal injury. Elife 11. 10.7554/eLife.76254

Wu, W., Nguyen, T., Ordaz, J.D., Zhang, Y., Liu, N.K., Hu, X., Liu, Y., Ping, X., Han, Q., Wu, X., Qu, W., Gao, S., Shields, C.B., Jin, X., Xu, X.M., 2022. Transhemispheric cortex remodeling promotes forelimb recovery after spinal cord injury. JCI Insight 7. 10.1172/JCI.INSIGHT.158150

Yang, L., Martin, J.H., 2023. Effects of motor cortex neuromodulation on the specificity of corticospinal tract spinal axon outgrowth and targeting in rats. Brain Stimul 16, 759–771. 10.1016/J.BRS.2023.04.014

Yang, Q., Ramamurthy, A., Lall, S., Santos, J., Ratnadurai-Giridharan, S., Zareen, N., Alexander, H., Ryan, D., Martin, J.H., Carmel, J.B., 2019. Independent replication of motor cortex and cervical spinal cord electrical stimulation to promote forelimb motor function after spinal cord injury in rats. Exp Neurol 320, 112962. 10.1016/J.EXPNEUROL.2019.112962

Yang, Y., Chen, X., Yang, C., Liu, M., Huang, Q., Yang, L., Wang, Y., Feng, H., Gao, Z., Chen, T., 2024. Chemogenetic stimulation of intact corticospinal tract during rehabilitative training promotes circuit rewiring and functional recovery after stroke. Exp Neurol 371, 114603. 10.1016/J.EXPNEUROL.2023.114603

Zheng, B., Tuszynski, M.H., 2023. Regulation of axonal regeneration after mammalian spinal cord injury. Nat Rev Mol Cell Biol 24, 396–413. 10.1038/S41580-022-00562-Y

