## Supplemental Figure 1 for "Chronic activation of corticospinal tract neurons after pyramidotomy injury enhances neither behavioral recovery nor axonal sprouting"

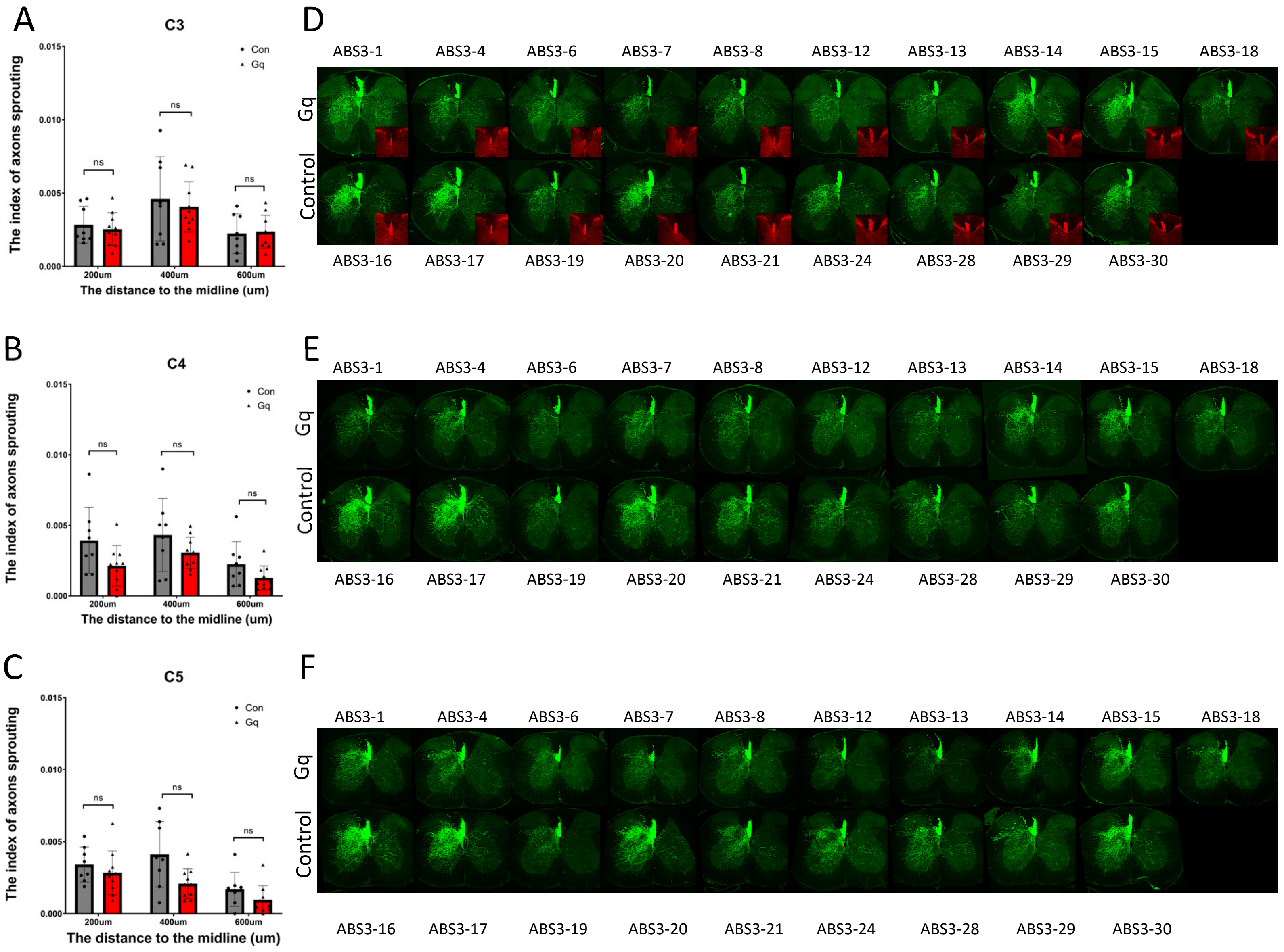


**Supplementary Figure 1. Images of all analyzed animals confirm consistent pyramidotomy injury and similar levels of cross-midline sprouting in cervical spinal cord.** (A,B,C) show quantification of cross-midline CST sprouting at individual cervical levels C3 (A), C4 (B), and C5 (C). (D,E,F) show images of transverse sections of cervical spinal cord with CST axons labeled by EGFP (green). Insets in (D) show PCKγ immunohistochemistry for each animal, confirming unilateral ablation of the corticospinal tract. ns indicates p>.05, 2-way ANOVA with Tukey's post hoc test. Error bars show ±SEM.
